# Histone acetylation at the sulfotransferase 1a1 gene is associated with its hepatic expression in normal aging

**DOI:** 10.1101/2020.09.17.300657

**Authors:** Mohamad M. Kronfol, Mikhail G. Dozmorov, Fay M. Jahr, Matthew S. Halquist, MaryPeace McRae, Dayanjan S. Wijesinghe, Elvin T. Price, Patricia W. Slattum, Joseph L. McClay

## Abstract

Phase II drug metabolism is poorly studied in advanced age and older adults may exhibit significant variability in their expression of phase II enzymes. We hypothesized that age-related changes to epigenetic regulation of genes involved in phase II drug metabolism may contribute to these effects. We examined published epigenome-wide studies of human blood and identified the *SULT1A1* and *UGT1A6* genes as the top loci showing epigenetic changes with age. To assess possible functional alterations with age in the liver, we assayed DNA methylation (5mC) and histone acetylation changes around the mouse homologs *Sult1a1* and *Ugt1a6* in liver tissue from mice aged 4-32 months obtained from the National Institute on Aging rodent tissue bank. Our sample shows significant loss of 5mC at *Sult1a1*, mirroring the loss of 5mC with age observed in human blood DNA at the same locus. We also detected increased histone 3 lysine 9 acetylation (H3K9ac) with age at *Sult1a1*, but no change to histone 3 lysine 27 acetylation (H3K27ac). *Sult1a1* gene expression is significantly positively associated with H3K9ac levels, accounting for 23% of the variation in expression. We did not detect any significant changes at *Ugt1a6*. We conclude that *Sult1a1* expression is under epigenetic influence in normal aging and that this influence is more pronounced for H3K9ac than DNA methylation or H3K27ac in this study. More generally, our findings support the relevance of epigenetics in regulating key drug metabolizing pathways. In future, epigenetic biomarkers could prove useful to inform dosing in older adults.

## Introduction

Age-associated changes to hepatic phase II drug metabolism is poorly understood. Recently, it has been suggested that epigenetic mechanisms could regulate genes encoding drug metabolizing enzymes in older adults^1^. Aging is known to be associated with extensive changes to epigenetic marks such as 5-methylcytosine (5mC)^2^ and, crucially, these changes are not purely stochastic. Several hundred loci in the genome exhibit consistent 5mC changes in normal aging and these are known as age-associated differentially methylated regions (a-DMRs). Many a-DMRs have functional consequences^2,3^, leading us to seek out a-DMRs at genes encoding drug metabolizing enzymes to determine if they affect regulation of these genes. Previously, we reported that a-DMRs at the cytochrome P450 2E1 gene, which encodes a phase I drug metabolizing enzyme, showed significant changes with age that were associated with CYP2E1 function^4^. In the current study, we extend this work to consider genes encoding phase II (conjugation) drug metabolizing enzymes.

Data from epigenome-wide association studies (EWAS) in human blood have shown significant changes to 5mC at several phase II genes with age^5–8^. However, it is unknown if these a-DMRs are present and have functional effects on gene expression in the liver, the primary organ of drug metabolism. Therefore, our goal in this study is to identify age-related changes to 5mC and histone acetylation at genes encoding phase II drug metabolizing enzymes and test if these epigenetic changes are associated with hepatic gene expression. This study used liver tissue from mice aged under controlled conditions, obtained from the National Institute on Aging tissue bank. This mouse model affords high experimental control and availability of relevant tissue.

We identified two phase II drug metabolism genes, *SULT1A1* and *UGT1A6*, showing evidence for a-DMRs in human blood studies (Supplementary **Table S1**). We mapped the associated sites in the human genome to their homologous mouse regions and tested for epigenetic changes at these regions in mouse liver (**Figure 1**). The mouse regions investigated in this study are highly conserved and show clear homology with their human counterparts^9,10^. High-Resolution Melt (HRM) analysis of bisulfite-converted DNA and chromatin immunoprecipitation (ChIP) assays were used to measure 5mC and histone post-translational modifications respectively^11^. We observed significant age-related epigenetic change at the *Sult1a1* gene and confirmed that this change was strongly associated with its hepatic gene expression.

**Figure 1.**
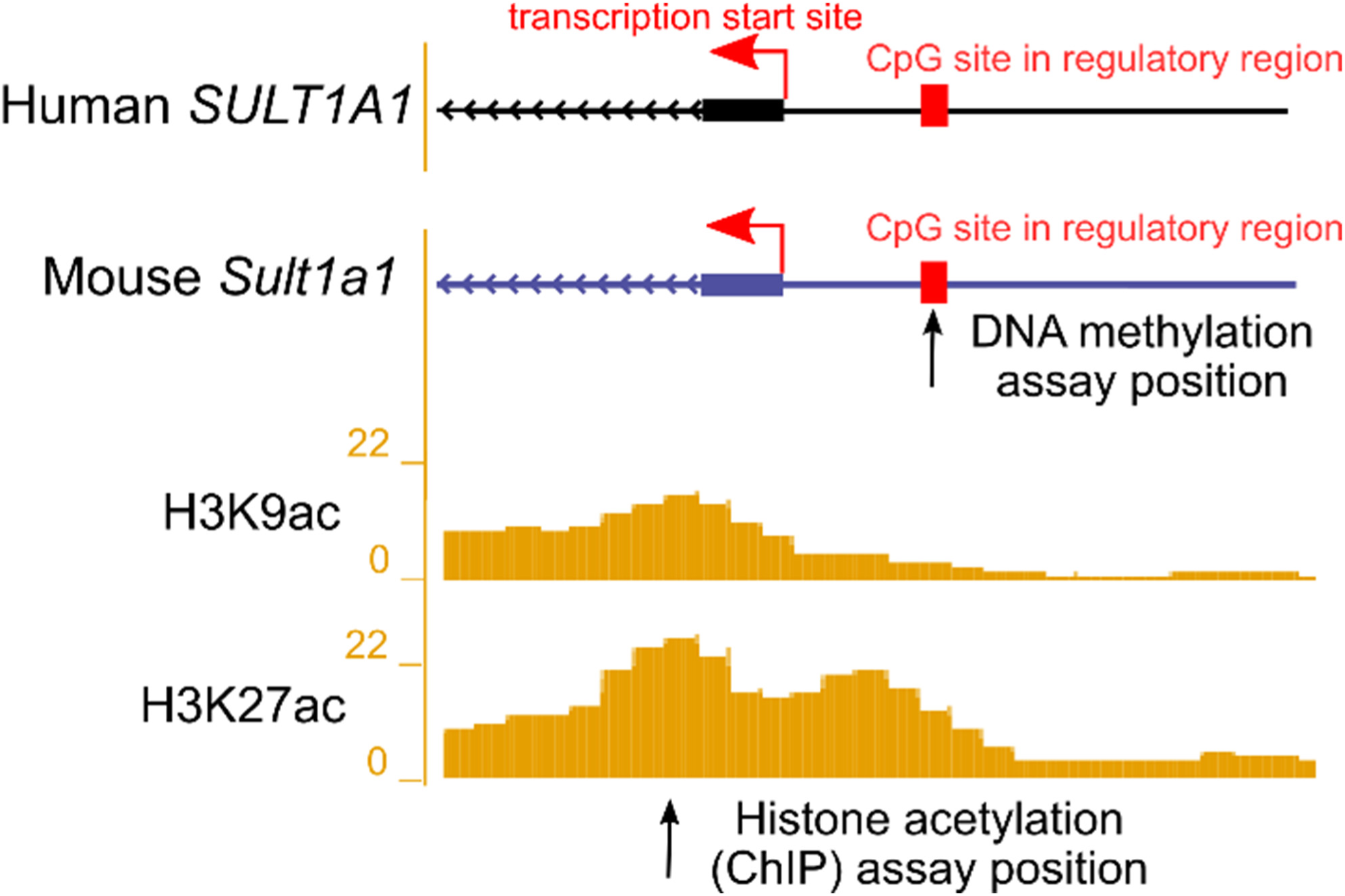
Illustration showing the transcription start site (TSS) and upstream regulatory regions of the human *SULT1A1* and the homologous mouse *Sult1a1* gene. DNA methylation assays were conducted in the current study at the upstream regulatory region in mouse at the position marked by the arrow. Reference histone 3 lysine 9 acetylation (H3K9ac) and histone 3 lysine 27 acetylation (H3K27ac) in 8-week-old mouse livers are from the ENCODE/LICR track in UCSC Genome Browser. A locus with high liver histone acetylation levels was chosen for analysis using ChIP-qPCR in the current study, at the position marked by the arrow. For exact assay coordinates, see Supplementary Table S2.

## Methods

Additional details are in the Supplementary Material.

### Samples

We obtained livers from 20 male CB6F1 mice from the National Institute on Aging (NIA) rodent tissue bank. Each of four age groups (4, 18, 24, and 32 months) comprised five subjects. We used the AllPrep DNA/RNA kit (Qiagen, Hilden, Germany) to extract DNA and RNA from the same liver samples. Quantity and purity of nucleic acids were measured using a Nanodrop spectrometer (ThermoFisher, Waltham, MA).

### Selection of genomic regions of interest

We obtained results for published EWAS of aging in human blood DNA (see Dozmorov 2015^11^) and contrasted these with lists of genes involved in ADME (absorption, distribution, metabolism, and excretion) processes from the pharmaADME consortium (pharmaADME.org) (Supplementary **Table S1**). Genomic coordinates of all regions investigated in mouse are provided in Supplementary **Table S2**. Using publicly available data (GEO accession numbers GSM1000153, GSM1000140), we identified regions around our target genes with high levels of histone 3 lysine 9 acetylation (H3K9ac) and histone 3 lysine 27 acetylation (H3K27ac) (**Figure 1**).

### Bisulfite conversion and High-Resolution Melt (HRM) analysis

200ng of liver genomic DNA per sample was treated with sodium bisulfite using the EZ DNA Methylation kit (Zymo Research, Irvine, CA). Standards using mouse genomic DNA of known percentage methylation (0, 5, 25, 50, 75, and 100% 5mC) were obtained from EpigenDx (Hopkinton, MA) and included on each assay plate. HRM assays using MeltDoctor reagents on a Quantstudio 3 (Applied Biosystems, Foster City, CA) were used to measure 5mC levels on the promoter and exon 2 of *Sult1a1* and *Ugt1a6* respectively (Supplementary **Table S2**). Samples were amplified by qPCR as follows: 10min hold at 95°C followed by 45 cycles of: 15sec at 95°C, 60sec at 60°C or 55°C for *Sult1a1* or *Ugt1a6* respectively, followed by a melt curve stage. Each reaction included 20ng bisulfite-converted DNA, 0.2μM forward and 0.2μM reverse primer (Supplementary **Table S2**). All DNA samples and standards were run in triplicate. Net Temperature Shift values^12^ of the liver DNA samples were interpolated on the standard curve to estimate their 5mC percentage.

### Gene expression analysis by reverse transcription – quantitative PCR (RT-qPCR)

Gene expression was measured for *Sult1a1, Ugt1a6a* and *Ugt1a6b*. We measured expression of both isoforms of *Ugt1a6* present in mouse liver because they are both expressed from the same locus and the regulatory region investigated could affect either of them. For each sample, 1μg of total liver mRNA was reverse transcribed using the iScript kit (Bio-Rad, Hercules, CA). Aliquots of cDNA were amplified in triplicate using TaqMan master mix (Applied Biosystems) and TaqMan *Sult1a1* or *Ugt1a6a/b* Mouse Gene Expression Assay (Mm01132072_m1, Mm01967851_s1, Mm03032310_s1, ThermoFisher). qPCR used 2 min hold at 50 ^°^C then 10min at 95 °CC, followed by 40 cycles of 15 sec at 95°CC and 60 sec at 60°CC. Murine *Gapdh* expression was used as the endogenous control gene (Mm99999915_g1, ThermoFisher). Quantification cycles (C_q_) were determined using the Relative Quantification application on the ThermoFisher Cloud. Normalized quantification cycles (ΔC_q_) were calculated by subtracting mean *Gapdh* C_q_ from the mean C_q_ of any of the genes.

#### Chromatin Immunoprecipitation Quantitative Polymerase Chain Reaction (ChIP-qPCR)

The TruChIP tissue shearing kit (Covaris, Woburn, MA) was used to process 80 mg of mouse liver per sample. This involved initial fragmentation on dry ice, followed by fixation in 1% formaldehyde for 2 min, freezing in liquid N2, mechanical pulverization and cell lysis. Chromatin was sheared on a Covaris M220 for 8min. 2% of sheared chromatin per IP was retained as input control. Each ChIP used 2μg sheared chromatin and was incubated overnight (16 hours) at 4°C with either 5μg of anti-H3K9ac (39137, Active Motif, Carlsbad, CA), or 5μg of anti-H3K27ac (39133, Active Motif). A separate control ChIP was also performed for each sample using 5μg of Rabbit IgG (ab171870, Abcam). After antibody incubation, ChIP mixtures were added to Dynabeads Protein G (ThermoFisher) for 4 hours at 4°C before washing and elution at 65°C for 1 hour. ChIPs were incubated with RNAse followed by Proteinase K and DNA purified using QIAquick (Qiagen).

Each ChIP DNA sample was assayed in triplicate using PowerUp SYBR Green qPCR master mix (Applied Biosystems) and 0.2 μM of forward and reverse primers (see Supplementary **Table S2** for primer sequences). Cycling conditions were: 2min at 50°C, 2min at 95°C followed by 45 cycles of: 15sec at 95°C, 30sec at 58 °C or 60°C for *Sult1a1* or *Ugt1a6* respectively and 1min at 72°C, followed by melt curve stage. For each sample, the mean threshold cycle (Cq) was normalized to the dilution factor (2%=1/50) corrected Cq value (Log2 (50) =5.6438) of the input control to obtain ΔC_q_. Percentage of input was calculated by multiplying 100 by 2^ΔCq^.

### Statistics

Linear regression analysis was performed in R version 3.6.1 (www.r-project.org) with α=0.05.

## Results

### Age-associated changes to Sult1a1 and Ugt1a6 5mC and gene expression

We obtained an average 260/280 value of 1.94 [1.8-2.1] for DNA and 2.05 [1.94-2.1] for RNA, showing that our extractions from liver were successful. HRM analysis of genomic DNA revealed that 5mC decreases significantly with age at *Sult1a1* (β=-1.08, 95%CI [-1.8, −0.2], SE=0.38, p=0.011) (**Figure 2a**) but there was no significant change at *Ugt1a6* (**Supplementary Figure 1a**).

**Figure 2.**
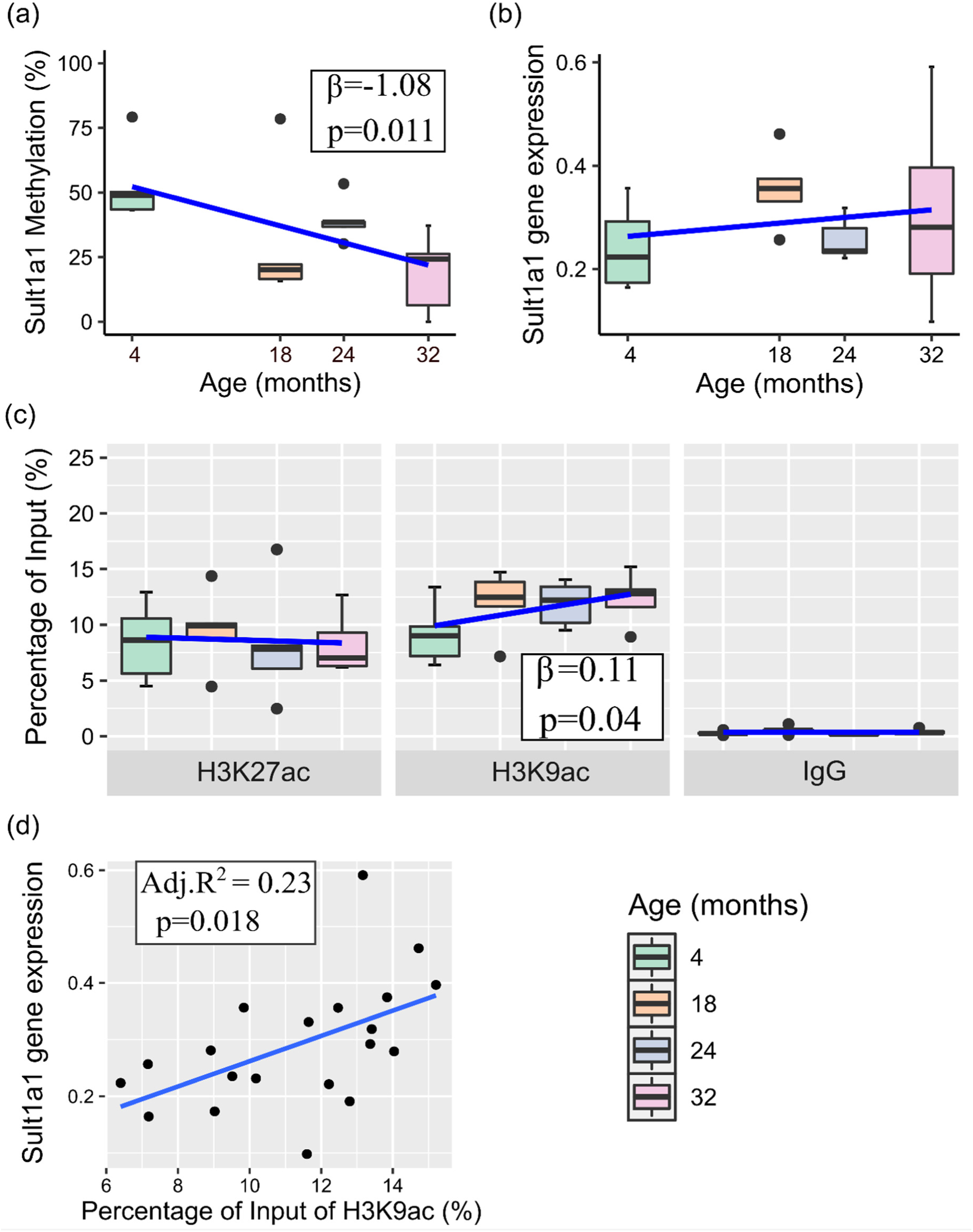
Box plots with superimposed regression line (blue) of Age-Associated changes to *Sult1a1* (a) methylation (n=20), (b) gene expression (n=20), and (c) Histone 3 Lysine 27 and Lysine 9 acetylation (H3K27ac and H3K9ac), chromatin Immunoprecipitation quantitative polymerase chain reaction (ChIP-qPCR) data (n=20 per target), IgG percentage of input shows a low background noise signal for each of the sample’s age groups. Data represent median (middle hinge), 25% (lower hinge) and 75% (upper hinge) quantile. Data points beyond upper or lower 1.5 * Inter Quantile Range are represented as individual black dots. (d) linear regression plot of percentage of input of H3K9ac and *Sult1a1* gene expression (n=20), with reported Adjusted R^2^ and p-value.

The observed *Sult1a1* promoter 5mC decrease with age translates to a 24% decrease in the 32 months group versus the 4 months group. On the other hand, neither of *Sult1a1* or *Ugt1a6a/b* gene expression changed in a consistent manner with chronological age in this sample (**Figure 2b, Supplementary Figure 1b and c)**.

### Histone acetylation analysis of Sult1a1 regulatory region

H3K9ac increased significantly with age at *Sult1a1* (β= 0.11, 95%CI [0.002,0.22], SE=0.05, p=0.04) **(Figure 2c)** but there was no significant change at *Ugt1a6* (Supplementary **Figure 1d**). The observed change with age at *Sult1a1* corresponds to a 0.11% increase in H3K9ac per month. H3K27ac levels did not change with age at either gene this sample **(Figure 2c**, Supplementary **Figure 1d)**

### Sult1a1 epigenetics and expression

As outlined above, 5mC and H3K9ac levels at *Sult1a1* changed with age. However, *Sult1a1* gene expression was not significantly associated with chronological age. To determine if epigenetic changes impacted *Sult1a1* gene expression, we tested the degree of association between 5mC and H3K9ac at *Sult1a1* with its gene expression. H3K9ac changes on *Sult1a1* intron1 are associated with its gene expression (β=0.02, 95%CI [0.004,0.04], SE=0.008, p=0.018), explaining 23% of the variability of the latter (Adj.R^2^=0.23, p=0.018) **(Figure 2d**). However, 5mC at the *Sult1a1* promoter was not associated with its gene expression (data not shown). This result suggests that H3K9ac is a more robust independent predictor of gene expression of *Sult1a1* than 5mC or chronological age.

## Discussion

This study demonstrates that 5mC and H3K9ac levels change with age at *Sult1a1* in mouse liver. Moreover, H3K9ac was associated with *Sult1a1* gene expression, while chronological age was not. This suggests that histone acetylation levels may be a better predictor of drug sulfonation by *Sult1a1* in advanced age than chronological age itself.

Our findings support prior observations in human blood studies. In particular, Reynolds *et al*. (2014)^7^ reported hypomethylation with age at human *SULT1A1*. We assayed the mouse genomic region homologous to the Reynolds *et al*. finding and showed that it exhibited hypomethylation with age in mouse liver too, thereby demonstrating consistency of the effect across tissues and species. Fu et al. (2012)^13^ reported increased *Sult1a1* expression in mouse liver tissue aged 27 months compared to 3 months. This suggests that increases to *Sult1a1* gene expression could occur with age but require larger sample sizes than used here to be reliably detectable. Age-related changes to H3K9ac levels were detectable at *Sult1a1* in our sample and H3K9ac is associated with its gene expression to a substantial degree of 23%, indicative of its functional importance as a determinant of gene activity. Prior studies of histone post-translational modification dynamics in aging are limited and we are not aware of existing data for *Sult1a1* in the liver. Schroeder et al. (2013)^14^ found that histone deacetylase inhibitors influenced gene expression of *Sult1a1* in the brain, but this study did not examine the effect of age nor changes in the liver.

Current understanding of phase II metabolism in old age is limited^15^ and sulfotransferase activity is poorly studied in older adults. There have been limited reports of decreased sulfonation in frail older adults for drugs such as metoclopramide^16^ but metoclopramide is primarily metabolized by SULT2A1^17^. To the best of our knowledge, the sulfonation of drugs by SULT1A1 in old age has not been investigated. SULT1A1 comprises the majority, 53%, of the total hepatic sulfotransferase^18^. SULT1A1 substrates include tamoxifen, levodopa, and estrogen replacement therapies^19,20^ and these drugs are indicated for the treatment of age-related disease or have detrimental and unpredictable toxicities. For example, SULT1A1 activates the prodrug tamoxifen which is essential for its anti-estrogen activity in the treatment of breast cancer ^21^. Levodopa, indicated for Parkinson’s disease in combination of carbidopa and entacapone, is primarily metabolized by SULT1A1 and SULT1A3^20^. Interestingly, prior studies have associated genetic sequence variants at *SULT1A1* with response to estrogen-replacement therapy^22,23^. Therefore, it is conceivable that age-related functional epigenetic changes at *SULT1A1*, as shown here, may similarly affect response to estrogen-replacement therapy. This area warrants future clinical study in humans.

A limitation of our study is that we could not directly manipulate epigenetic levels in our liver samples, so the relationship between epigenetics and gene expression is correlational, not causal. Nevertheless, the magnitude of the correlation between epigenetics and hepatic *Sult1a1* gene expression suggests that further study of pharmacoepigenetic biomarkers may be fruitful. Human pharmacokinetic studies should be undertaken to determine if epigenetic biomarkers could meaningfully predict rates of drug metabolism in older adults.

## Study Highlights

### What is the current knowledge on the topic?

The determinants of age-related changes of drug response due to phase II metabolism are poorly understood. The study of epigenetic regulation of ADME genes is in its infancy and no reports correlate age-related changes to epigenetic regulation of genes controlling phase II metabolism.

### What question did this study address?

What is the extent of age-related change of DNA methylation and histone acetylation at the *Sult1a1* and *Ugt1a6* genes*?* What is the extent of association of these epigenetic marks with gene expression?

### What does this study add to our knowledge?

We determined the extent of change to DNA methylation and histone acetylation on sulfonation gene, *Sult1a1*, and its degree of association with hepatic gene expression in advanced age. The effect of histone acetylation (H3K9ac) on expression was more pronounced than that of DNA methylation.

### How might this change clinical pharmacology or translational science?

Our findings could stimulate clinical work to determine how we dose older patients based on epigenetic marks at genes controlling phase II metabolism, to optimize drug response and decrease adverse events.

## Acknowledgements

MMK was partially funded through a graduate studentship from Virginia Commonwealth University (VCU) School of Pharmacy and this study was completed in partial fulfillment of the doctoral requirements in Pharmaceutical Sciences at VCU. We are grateful to the staff at the NIA Rodent Tissue Bank for providing us with the samples to carry out this study.

## Author Contributions

MMK designed research, performed research, analyzed data and wrote the manuscript MGD helped design the study, analyzed data and wrote the manuscript

FMJ performed research, provided laboratory support and wrote the manuscript

MSH, MPM, DSW, ETP and PWS helped design the study, offered critical feedback and wrote the manuscript

JLM designed the study, obtained funding, analyzed data and wrote the manuscript

## Supplementary Information Titles

In Supplement:

### Supplementary Methods

Table S1. Summary of human EWAS findings for phase II drug metabolism genes

Table S2. HRM and ChIP primers and qPCR product sequences

Figure S1. *Ugt1a6* epigenetic modifications and gene expression with age

