## Supplementary Material for "Histone acetylation at the sulfotransferase 1a1 gene is associated with its hepatic expression in normal aging"

### Supplementary Methods

*Bisulfite conversion of genomic DNA and High-Resolution Melt (HRM) analysis:*

200ng of liver genomic DNA was treated with sodium bisulfite according to the EZ DNA Methylation kit protocol (D5002, Zymo Research, Irvine, CA). Mouse genomic DNA of 0, 5, 25, 50, 75, and 100% methylation were used as standards (808060M, EpigenDx, Hopkinton, MA). A negative (no template) control was also used per plate. High-Resolution Melt (HRM) assays using MeltDoctor HRM MasterMix (4409535, Applied Biosystems, Foster City, CA) on a Quantstudio 3 instrument were used to measure 5mC levels at 134 and 206 base pairs regions on the promoter and exon 2 of mouse *Sult1a1* and *Ugt1a6* encompassing 1 and 4 CpGs respectively (chr7:133,820,382-133,820,515, chr1:90,035,169-90,035,374 mouse genome assembly mm9 NCBI37/ build 9, July 2007) (Supplementary Table S2). Samples were amplified by qPCR as follows 10min hold at 95ºC followed by 45 cycles of: 15sec at 95ºC, 60sec at 60ºC or 55ºC for *Sult1a1* or *Ugt1a6* respectively, followed by a melt curve stage with temperature range of 60ºC to 95ºC with fluorescence capture at 0.15 degrees per second increment. Each reaction included 20ng bisulfite-converted DNA and final concentration of 1X MeltDoctor HRM MasterMix (Applied Biosystems), 0.2µM forward and 0.2µM reverse primer (Supplementary Table S2). The liver samples and the standards were run in triplicate. The Net Temperature Shift (NTS) values of the liver samples were interpolated on the standard curve to yield their 5mC percentage.

*Gene expression analysis by reverse transcription – quantitative PCR (RT-qPCR):*

For each sample, 1µg of total liver mRNA was reverse transcribed to cDNA by iScript kit according to manufacturer's protocol (1708891, Bio-Rad, Hercules, CA). Aliquots of cDNA were amplified in triplicate using final concentration of 1X TaqMan master mix (4369016, Applied Biosystems) and 1X TaqMan *Sult1a1* or *Ugt1a6a/b* Mouse Gene Expression Assay (Mm01132072_m1, Mm01967851_s1, Mm03032310_s1 ThermoFisher) [11–13]. qPCR conditions were as follows 2 min hold at 50 °C then 10min at 95 °C, followed by 40 cycles of 15 sec at 95°C and 60 sec at 60°C. Mouse *Gapdh* endogenous control was also assayed in triplicate (Mm99999915_g1, ThermoFisher) [14–16]. A 1:2 serial dilution of control cDNA was used to generate a 5-point standard curve. Quantification cycles ($C_q$) were determined using the Relative Quantification application on the ThermoFisher Cloud. Normalized quantification cycles ($\Delta C_q$) were obtained by subtracting the mean *Gapdh* $C_q$ from the mean *Cyp2e1* $C_{q.}$

*Chromatin Immunoprecipitation (ChIP) – qPCR:*

80 mg of mouse liver was minced into 1 mm3 fragments with -20 C precooled scalpel and transferred to 2mL Eppendorf tube. TruChIP tissue shearing kit (520237, Covaris, Woburn, MA) was used to prepare sheared chromatin. Centrifugations were at 200g for 5 min at 4ºC unless otherwise noted. The minced tissue was washed with 1 mL 1X cold PBS and centrifuged at 200g for 5min at 4ºC. The supernatant was discarded and 1mL of fixing buffer A was used to resuspend the pellet. 100µL of freshly prepared 11.1% methanol-free formaldehyde (1% final concentration) was added and the fixation was quenched after 2 min with 58µL quenching buffer E. The suspension was centrifuged, and the supernatant discarded. Pellet was washed twice with 1mL 1X cold PBS and centrifuged. Pellet was transferred to tissueTUBE (TT05M XT) provided in the kit and flash frozen in liquid nitrogen for 45sec and pulverized into powder by a precooled pestle. Pulverized tissue was transferred by inverting to a screwed-on milliTUBE-2mL and stored at -80ºC until nuclei separation and shearing step the next day. Pulverized tissue was transferred to a 2mL Eppendorf tube using two successive transfers by 500µL lysis buffer B. The 2mL tube was incubated on a rotor at 4ºC for 20min to complete lysis. Nuclei were pelleted by centrifugation at 1700g for 5min at 4ºC. Supernatant was discarded and pellet was resuspended in 1mL wash buffer C and incubated on a rocker for 10min at 4ºC. Nuclei were pelleted again at 1700g for 5min at 4ºC and supernatant discarded. Pellet was resuspended with 1mL wash buffer C and centrifuged at 1700g for 5min at 4ºC. Pellet was resuspended with 1mL shearing buffer D2 and sheared on M220 (Covaris, Woburn, MA) for 8min at 75 PIP, 10% duty factor, 200CPB, 7C set point temperature (4/10 Min/Max), and no degassing. Time course trials were conducted to determine optimal shearing (2-20 min) and fixation times (2 and 5 min) of 8 and 2 min respectively which yielded the highest percentage (>75%) of fragment sizes between 150 and 700 bp and the lowest percentages (<25%) of fragment sizes less than 150 and

higher than 701 bp combined. Shearing and fixation time course trials were analyzed on the Agilent 2100 Bioanalyzer using the Agilent DNA 12000 chip on Agilent 2100 expert software. 25µL of sheared chromatin was incubated with 1µL 10mg/ml RNAse (EN0531, Thermo Fisher, Waltham, MA) at 37ºC for 30min, then it was treated with 4µL 10mg/ml Proteinase K (17916, Thermo Fisher, Waltham, MA) at 65ºC overnight (16 hours). DNA was purified using QIAquick PCR purification Kit (Qiagen, Hilden, Germany). The concentration of eluted DNA was measured on Qubit 4.0 fluorometer and used to calculate the volume required to have 2µg sheared chromatin as starting material for the Chromatin immunoprecipitation (ChIP) step (1:1 ratio of DNA to chromatin was used to calculate chromatin concentration). The sheared chromatin was diluted 1:2 with 3X Covaris IP dilution buffer in order to decrease final SDS concentration to 0.083% and prevent SDS interference with epitope and antibody binding.

Each ChIP had 2µg of sheared chromatin. 2% of the volumes of sheared chromatin per IP was set aside at 4ºC as input control and was not processed through ChIP step. ChIP was carried out by incubating 5 µL of H3K9ac (39137, Active Motif, Carlsbad, CA), or 5µL of H3K27ac (39133, Active Motif, Carlsbad, CA), or 5µL of Rabbit IgG (ab171870, Abcam) with 2µg sheared chromatin from each sample overnight (16 hours) at 4ºC. Hence, three worth of ChIP volumes were used from each sample. The formed complex was incubated with 50 µL Dynabeads Protein G (10003D, Thermo Fisher, Waltham, MA) for 4 hours at 4ºC. The bead linked complex was inserted on the DynaMag (Thermo Fisher, Waltham, MA) magnet rack for 2 min and the supernatant discarded. The bead coupled complex was removed from the magnet and washed 3 times with 500 µL with cold 0.05X Tween 20 in PBS PH7.4 (10010023, Thermo Fisher, Waltham, MA) for 3 min each at room temperature on HulaMixer (Thermo Fisher, Waltham, MA). Finally, 50 µL of IP elution buffer (Aq. 1% SDS, 0.1 M NaHCO3, pH 9.0) was added to the bead coupled complex and incubated on a heat block for 1hour at 65ºC with 15sec vortexing every 15 min. The samples were then inserted back on the DynaMag for 2 min and 50 µL of the supernatant was transferred to a 96 well PCR plate. The input control was diluted 1:2 with 3X Covaris IP dilution buffer to mimic the dilution done to the samples and preserve its percentage (2%). The ChIP'ed samples and their input controls were incubated each with 2 µL RNAse for 30 min at 37C and then 8 µL of Proteinase K was added and incubated overnight (16 hours) at 65 ºC. The next day, DNA was purified using QIAquick PCR purification kit (Qiagen, Hilden, Germany) to final elution volume of 50 µL. 2 µL of each DNA elution (H3K9ac, H3K27ac, IgG, and input control) per sample was used per qPCR reaction.

In each qPCR reaction, 2 µL of DNA was amplified in triplicate on the Quant Studio 3 instrument (Applied Biosystems). Reaction volumes was 20µL containing final concentration of 1X of PowerUp SYBR Green Master Mix (A25742, Applied Biosystems) and 0.2 µM of forward (0.4 µL) and reverse (0.4 µL) primers and 7.2 µL molecular biology grade H2O. The qPCR run conditions are as follows: 2min hold at 50 ºC followed by 2min hold at 95 ºC followed by 40 cycles of the following: 15sec at 95 ºC followed by 30sec at annealing temp at 58 ºC or 60ºC for *Sult1a1* or *Ugt1a6* respectively and by 1min at 72 ºC, followed melt curve analysis stage. This stage is as follows: 15sec hold at 95 ºC followed by 1min hold at 59 ºC and then the continuous fluorescence acquisition starts while increasing the temp by 0.15 ºC /s to end at 95 ºC followed by a 15sec hold at 95 ºC. Primer specificity and efficiency were tested on 2% agarose gel and a 5-point 1:2 serial dilution standard curve respectively. No secondary amplification or primer dimers were detected indicating primer specificity. A singular melt peak was shown by the qPCR melt curves indicating formation of a single product. Primer efficiency outside of accepted ranges [90-110] were discarded until reaction conditions were optimized. The Cq values for each plate were downloaded from ThermoFisher Cloud. The mean threshold cycle (Cq) values were obtained for each sample and normalized to the dilution factor (2% = 1/50) corrected Cq value (Log2 (50) =5.6438) of the input control to obtain Delta Cq. Percentage of input was calculated by multiplying 100 with 2 raised to the exponent of Delta Cq. For each sample, three "percentage of input" values were obtained and are H3K9ac, H3K7ac, and IgG.

**Table S1.** Summary of human EWAS findings for phase II drug metabolism genes

| | Longevity Map [1] | Heyn [2] | Reynolds [3] | Martilla [4] | Steegenga [5] | |
|---|---|---|---|---|---|---|
| | Exp | Meth | Meth | Both | Both | Total |
| ***SULT1A1*** | X | | X | | X | 3 |
| ***UGT1A6*** | | X | | X | | 2 |
| ***UGT2B15*** | | | | | X | 1 |
| ***GSTT1*** | X | | | | | 1 |
| ***SULT2B1*** | | | | | X | 1 |
| ***UGT1A4*** | | | | X | | 1 |
| ***UGT1A5*** | | | | X | | 1 |
| ***UGT1A1*** | | | | | | 0 |
| ***SULT1A3*** | | | | | | 0 |
| ***SULT1B1*** | | | | | | 0 |

Human ADME genes list from pharmaADME (www.pharmaadme.org) was contrasted on the top findings from EWAS and gene expression studies of normal aging in human blood DNA. Genes encoding phase II drug metabolizing enzymes that showed significant association in the top findings of these studies is marked with an "X". The total number of studies were a specific gene is a top association in the reported results is shown as a total in the last column of the table. The type of study is given as "Exp" for genome-wide gene expression, "Meth" for DNA methylation EWAS and "Both" for studies that encompassed both expression and DNA methylation.

**Table S2.** HRM and ChIP primers

| **Gene** | **Primers (5'→ 3')** | | **Product size/coordinates/assembly** | |
|---|---|---|---|---|
| | **Forward Primer** | **Reverse Primer** | | |
| | **High Resolution Melting** | | | **CpG count** |
| *Sult1a1* | GGAAGGTGTTTTTGTTTTTATG | CTAAAAATATATCTCTCCCAACT | 134/chr7:133,820,382-133,820,515/mm9 | 1 |
| *Ugt1a6* | AGTATGAAGGAGATAGTAGAATAT | AAAAAACCCATCAAAAAAACAC | 206/chr1:90,035,169-90,035,374/mm9 | 4 |
| | **Chromatin Immunoprecipitation quantitative Polymerase Chain Reaction** | | | |
| *Sult1a1* | GGGGAACTCAGACAAACCAC | TCCTGCCCAGATACTGGTTC | 154/ chr7:133,819,634-133,819,787/mm9 | |
| *Ugt1a6* | GCCACTCAGGAAGGACAGAG | GTGAGGCACTGGTCTGGTTT | 153/ chr1:90,030,630-90,030,782/mm9 | |

Primers used in the High-Resolution Melting (HRM) and Chromatin Immunoprecipitation quantitative Polymerase Chain Reaction (ChIP-qPCR) assays with size of PCR product, coordinates, and assembly with CpG count when appropriate.

**Figure S1** Age-associated changes to *Ugt1a6* epigenetic modifications and gene expression.

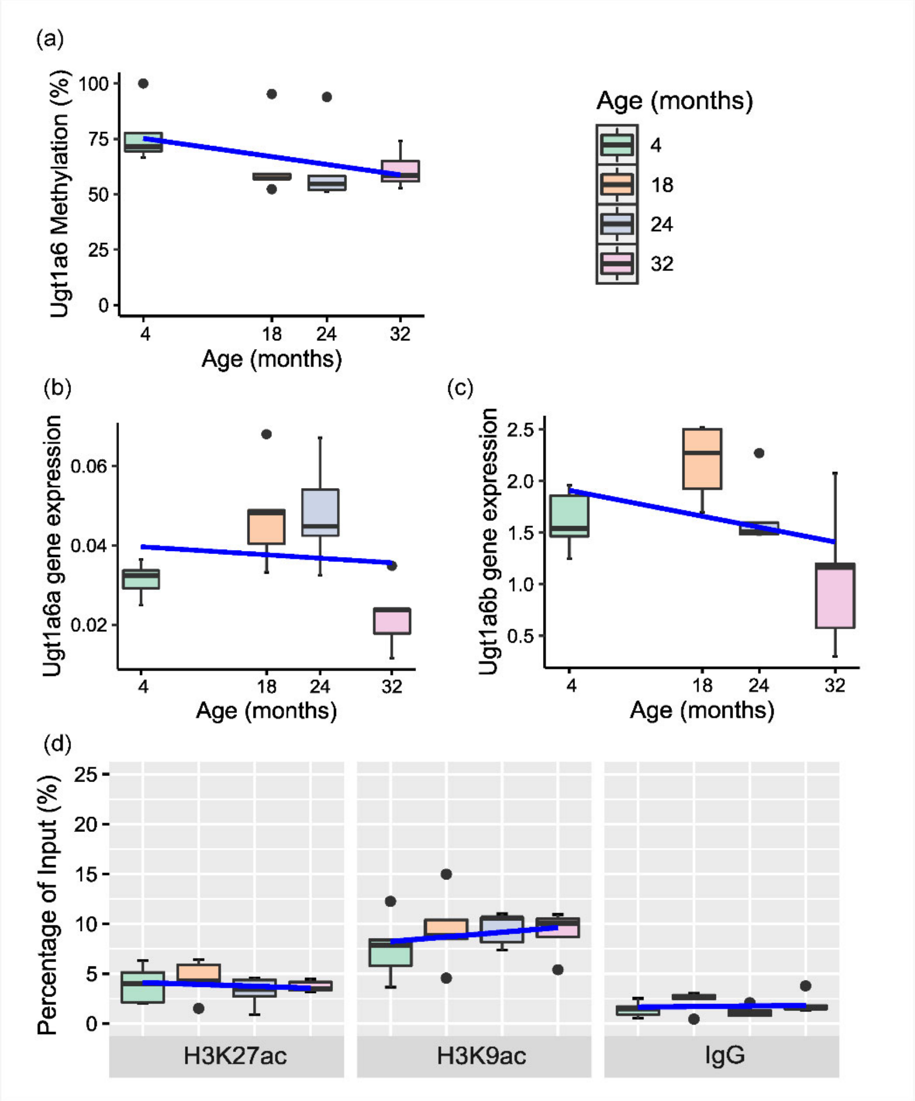


**Figure S1** Age-associated changes to *Ugt1a6* epigenetic modifications and gene expression. (a) Box plot with superimposed regression line (blue) of age-associated changes to *Ugt1a6* methylation (n=20) (p>0.05). Box plot with superimposed regression line (blue) of age-associated changes to gene expression of (b) *Ugt1a6a* isoform and (c) *Ugt1a6b* isoform (n=20 per locus) (p>0.05). (d) Histone 3 Lysine 27 and Lysine 9 acetylation (H3K27ac and H3K9ac), Chromatin Immunoprecipitation quantitative polymerase chain reaction (ChIP-qPCR) data (n=20 per target) (p>0.05), IgG percentage of input shows a low background noise signal for each of the sample's age groups. Data represent median (middle hinge), 25% (lower hinge) and 75% (upper hinge) quantile. Data points beyond upper or lower 1.5 * Inter Quantile Range are represented as individual black dots.
